# Decoding Graded Grip-Force Intensity from fMRI Data Reveals a Transformation from Abstract to Effector- and Movement-Specific Codes prior to Execution

**DOI:** 10.64898/2026.05.28.728406

**Authors:** Guido Caccialupi, Timo Torsten Schmidt, Felix Blankenburg

## Abstract

Motor planning entails a progressive transformation of neural representations—from abstract motor goals, which represent intended action-outcomes independent of any particular effector (i.e., the body part executing the action), to effector-specific movement plans. Functional MRI (fMRI) studies have shown that parametric variations in parietal activity patterns reflect the encoding of intended force intensities in effector-specific regions, even before detailed movement parameters are specified. However, how these intended force intensities are initially represented in an abstract, effector-independent format and subsequently transformed into effector- and movement-specific plans remains unclear. To address this, human participants performed a delayed grip-force task during fMRI. They first prepared two of four possible force intensities, then received a cue indicating which hand should apply which force, and finally executed both grips simultaneously. Using time-resolved support vector regression (SVR) combined with a searchlight approach, we identified brain regions that parametrically code grip-force intensities across two 6-second delay periods. During the first delay, above-chance decoding was observed in the precuneus (PCu), whereas during the second delay it emerged in effector-specific regions, including the contralateral intraparietal sulcus (pIPS/aIPS), primary somatosensory cortex (S1), dorsal premotor cortex (PMd), and supplementary motor area (SMA). Cross-decoding confirmed effector-independent coding in the PCu, while cross-temporal generalization revealed stable representations in the contralateral IPS and PMd from the second delay through execution. Together, these findings indicate a progressive transformation from abstract representations of intended force intensity in the PCu to effector- and movement-specific plans in the IPS and PMd.

## 1 INTRODUCTION

Motor planning is a complex neural process that underpins the ability to anticipate, represent, and specify parameters of upcoming actions prior to execution. It involves selecting the appropriate effector (i.e., the body part used to perform an action; Gallivan et al., 2011), maintaining relevant motor parameters in working memory (WM), and transforming abstract motor goals (i.e., representations of the intended action outcome prior to specification of effector or movement details; Majdandžić et al., 2007; Gallivan et al., 2011) into executable movement plans (Boettcher et al., 2021). In essence, motor planning can be interpreted as a sequence of processes that unfold over time: first, the intended abstract goal is represented; next, the effector is selected; and finally, the detailed movement required to achieve the goal is specified. Neuroimaging research has extensively investigated the neural correlates of goal encoding (Soon et al., 2008, 2013) and effector selection (Heed, 2011, 2016; Leoné et al., 2014), providing evidence that motor goals can be represented in effector-specific brain regions even before movement details are specified (Seegelke & Heed, 2025; Caccialupi et al., 2025b). However, it remains unclear how motor goals are represented in an abstract format prior to effector selection, and how these representations are transformed into effector-specific movement plans (Kim et al., 2021).

Foundational neurophysiological studies in non-human primates identified preparatory signals reflecting planned movements in the dorsal premotor cortex (PMd; Cisek & Kalaska, 2004, 2010; Hoshi & Tanji, 2006), supplementary motor area (SMA; Hoshi & Tanji, 2004), and posterior parietal cortex (PPC; Cui & Andersen, 2007, 2011; Andersen & Cui, 2009). Complementary neuroimaging studies in humans, using delayed-response paradigms (e.g., Hamilton & Grafton, 2003), have consistently shown that secondary motor areas, particularly the PMd, are involved in specifying movement features of grasping actions. For instance, univariate fMRI analyses in delayed grip-force tasks have shown that the BOLD signal amplitude in the PMd and intraparietal sulcus (IPS) covaries with anticipated grip-force intensity, indicating parametric coding of motor parameters prior to execution (Cole & Rotella, 2002; Chouinard et al., 2005; Nowak et al., 2009; van Nuenen et al., 2012; Mizuguchi et al., 2014). More recent fMRI multivoxel pattern analysis (MVPA) studies have further shown that above-chance decoding in parietal and premotor cortices can predict movement properties, such as movement sequences (Yokoi & Diedrichsen, 2019; Ariani et al., 2021, 2022), kinematics of grip types (Gallivan et al., 2011b; Ariani et al., 2015; Ruiz et al., 2024), and anticipated grip-force intensities (Caccialupi et al., 2025a, b). These findings corroborate the involvement of parieto-frontal networks in representing movement properties and anticipating detailed motor parameters, consistent with the final stage of motor planning.

Building on these findings, recent fMRI MVPA studies investigated where in the brain more abstract action features can be decoded independently of motor movements (Gallivan et al., 2011b; Ricciardi et al., 2013; Errante et al., 2021). In particular, information about intended action outcomes, or “motor goals”, has been decoded from PPC before movement preparation (Cattaneo et al., 2009; Gallivan & Culham, 2015; Turella et al., 2020; Liu et al., 2024; Gallivan et al., 2013; Ariani et al., 2018). These findings align with a two-stage model of motor planning in which abstract goals are first represented and then transformed into movement-specific plans (Wong et al., 2015; Boettcher et al., 2021). Consistently, the IPS has been recently found to contribute to the transformation of neural codes underlying the selection of intended actions into motor codes during an early planning phase (Caccialupi et al., 2025a). While both effector-independent and effector-specific information has been decoded from the IPS (Ariani et al., 2015; Gallivan et al., 2016; Heed et al., 2016; Caccialupi et al., 2025a, 2025b), the anterior IPS (aIPS) has been found to primarily supports effector-specific planning for hand actions (Leoné et al., 2014), whereas more posterior and medial PPC regions, including posterior IPS (pIPS) and precuneus (PCu), encode intended goals in an effector- and movement-independent format (Bode & Haynes, 2009; Gallivan et al., 2011, 2013; Heed et al., 2016; Turella et al., 2020). Together, these results indicate that distinct PPC subsections may represent action features of a motor plan at different levels of abstraction (Turella et al., 2020), providing a neural substrate for transformation of abstract motor goals into detailed, effector-specific movement plans.

The encoding of abstract motor goals and their transformation into detailed movement plans likely rely on neural mechanisms similar to those involved in WM transformations, which have been extensively investigated using fMRI decoding techniques (Haynes & Rees, 2006; Norman et al., 2006). Time-resolved MVPA in delayed-response paradigms has been essential for testing how, where, and when information is represented across different task periods (Schmidt et al., 2017; Hebart et al., 2018). For example, in WM studies, time-resolved decoding has been employed to track transition processes from low-level sensory features to more abstract representational codes across visual, tactile, and auditory modalities (Schmidt et al., 2009, 2017, 2021; Wu et al., 2018; Hebart et al., 2018; Uluç et al., 2018, 2020). These studies further indicate that transformations into more abstract representations occur in higher-order sensory or multimodal cortices, depending on task demands (Christophel et al., 2017). Recently, we extended the use of time-resolved MVPA to decode action-related representations, tracking the transformation processes underlying the selection of intended actions compared to detailed movement planning (Caccialupi et al., 2025a). In particular, using a delayed grip-force task, we found that intended grip-force intensities were initially represented in the ventromedial prefrontal cortex (vmPFC) and later transformed into motor codes in the l-PMd, revealing distributed neural patterns encoding grip-force parameters during grip-force selection and movement planning. In a subsequent study (Caccialupi et al. 2025b) we showed that contralateral IPS encodes intended grip-force intensities in an effector-specific format during an early planning stage. However, since effector information was always available to participants, these studies did not directly address where abstract motor goals (i.e., intended grip-force intensities) are represented before effector selection, nor how they are subsequently transformed into effector- and movement-specific plans.

The present study therefore investigates where and when abstract and effector-specific representations of grip-force intensities can be parametrically decoded from fMRI data. We modified our previous paradigm (Caccialupi et al., 2025a, 2025b) and employed a delayed grip-force task in which participants first maintained two of four possible force intensities, then selected which hand to prepare for which force, followed by bimanual execution. Using time-resolved MVPA with support vector regression (SVR), we decoded parametric representations of grip-force intensity during two subsequent delay periods. In the first 6-second delay, participants maintained the to-be-applied force intensities without knowing which hand would be used for which force. In the second delay, effector information was specified in preparation for execution. Additionally, cross-decoding analyses assessed whether grip-force intensities were represented in an effector-specific or effector-independent format, and temporal generalization tested whether multivariate activation patterns during execution were already present during the second delay, reflecting a movement-specific format during motor planning. Based on previous literature, we hypothesize that above-chance decoding will be initially observed in medial PPC regions, followed by lateralized action-specific regions, including the contralateral PMd and IPS.

## 2 MATERIALS AND METHODS

### 2.1 Participants

All participants (*N* = 31) were healthy and right-handed, as assessed by the *Edinburgh Handedness Inventory* (*EHI;* Oldfield, 1971). Before the experiment, they provided written informed consent to participate in the study in accordance with protocols approved by the local ethics committee of *Freie Universität Berlin* (003/2021, Berlin, Germany).

To account for potential biases and confounds, only *N* = 22 participants were included in the analysis (age: 28.9 ± 5.53, 11 female). Three participants were excluded due to excessive head movement (>3 mm), and six were excluded for frequent grip force application during at least one delay period (≥75% of trials in at least one run for condition). To further ensure that the analyses were not confounded by early motor responses, we excluded from the dataset all trials in which the grip force was applied during any delay period (cut-off: mean grip-force ≥ 0.05 on a scale from 0 to 1).

### 2.2 Experimental procedure

Before entering the MRI scanner, participants underwent a training session lasting 60 to 90 minutes approximately. Initially, the grip-force transducers were calibrated to participants’ maximum grip-force (i.e., the average of the maximum force applied by the right and left hands). Next, participants were introduced to the four grip-force levels that served as target grip-force intensities in the main experiment. In this training phase, they were familiarized with the transducer by performing sixteen trials with real-time visual feedback of the applied grip-force. In forty-eight further trials, they were trained on the experimental task (see below). Next, participants entered the MRI scanner, and the grip-force transducers were calibrated anew. During the acquisition of a structural MRI scan, participants were further trained on the experimental task. Finally, they performed the delayed grip-force task (illustrated in **Figure 1A**) in four runs during fMRI scanning.

**Figure 1.**
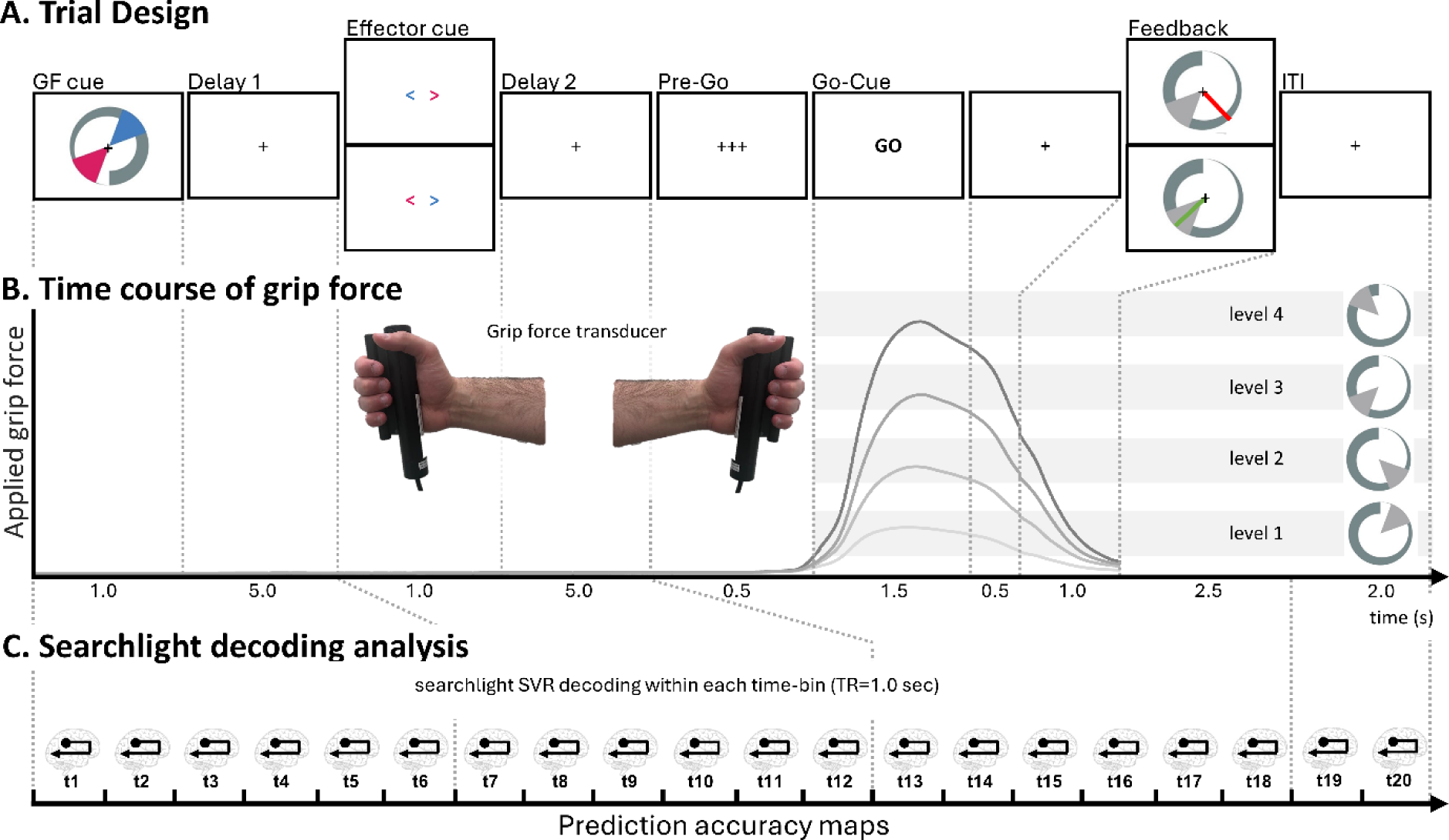
**A**. Delayed grip-force paradigm. Two grip-force levels were presented on a 1-second *Grip-force (GF) cue* (i.e., a grey circular increasing bar with one level represented in cyan and the other in red, as sectors of 60°) and maintained throughout a 5 s *Delay 1*. Next, a visual *Effector cue* (i.e., a pair of left- and right pointers) indicated what hand should be prepared for which force (based on the pointers orientation and colour) during a subsequent 5 s *delay 2*. After this second delay period, participants performed the task upon the presentation of a *pre-Go* (0.5 s) and a *Go-cue* (1.5 s). Subjects responded with their right and left hands by squeezing two grip-force devices with the cued intensities, and a *Feedback* representing the applied forces was provided at 2 s from the *Go-cue*. **B**. Lines in different shades of grey represent time courses of mean grip forces applied during the experimental-trial, calculated over the trials per condition (i.e., per grip-force level), normalized to the subjective maximum force, and averaged across hands. Light-grey bars displayed in the background represent grip-force levels. **C**. Representation of the multivoxel pattern analysis (MVPA) with a searchlight and a time-resolved approach adopted across three time periods of the trial, including: a 6 s *first delay period*, 6 s *second delay period*, and a 6 s *motor execution period*, *feedback* and *ITI*.

### 2.3 Grip-force assessment and stimuli

Based on the visual presentation of a *Grip-force cue* and an *Effector cue*, participants were instructed to simultaneously apply two different grip forces bimanually, one with the right hand and one with the left, using two cylindrical MR-compatible grip-force transducers (*i.e.*, *force fibre optic response pads*, Current Designs, HHSC-1x1-GRFC-V2; illustrated in **Figure 1B**). Grip-force intensity was sampled throughout the task to ensure that participants only applied force during the execution period of the trials, allowing the exclusion of trials where participants applied force during other periods of the experiment.

We used a well-established cuing procedure for the grip forces (see Caccialupi et al., 2025), based on the presentation of a visual *Grip-force cue* in the centre of the screen, utilizing *Psychtoolbox-3* (Brainard, 1997). The visual cue was composed of a 360° circular display increasing in thickness, corresponding to an increase in grip-force where the maximum corresponded to 75% of the individual maximum grip-force. Four grip-force levels were indicated by the display of a range, defining a sector of 60° of the grip-force spectrum, with grip-force level 1: 15°-75° (4.15-20.85% of maximum grip force level), level 2: 105°-165° (29.15-45.85%), level 3: 195°-255° (54.15-70.85%), and level 4: 285°-345°(79.15-95.85%) (as illustrated in **Figure 1B**, right side; and **Figure 2A**). The *Grip-force cue* was presented for each trial with a random degree of rotation (as illustrated in **Figure 1A**).

**Figure 2.**
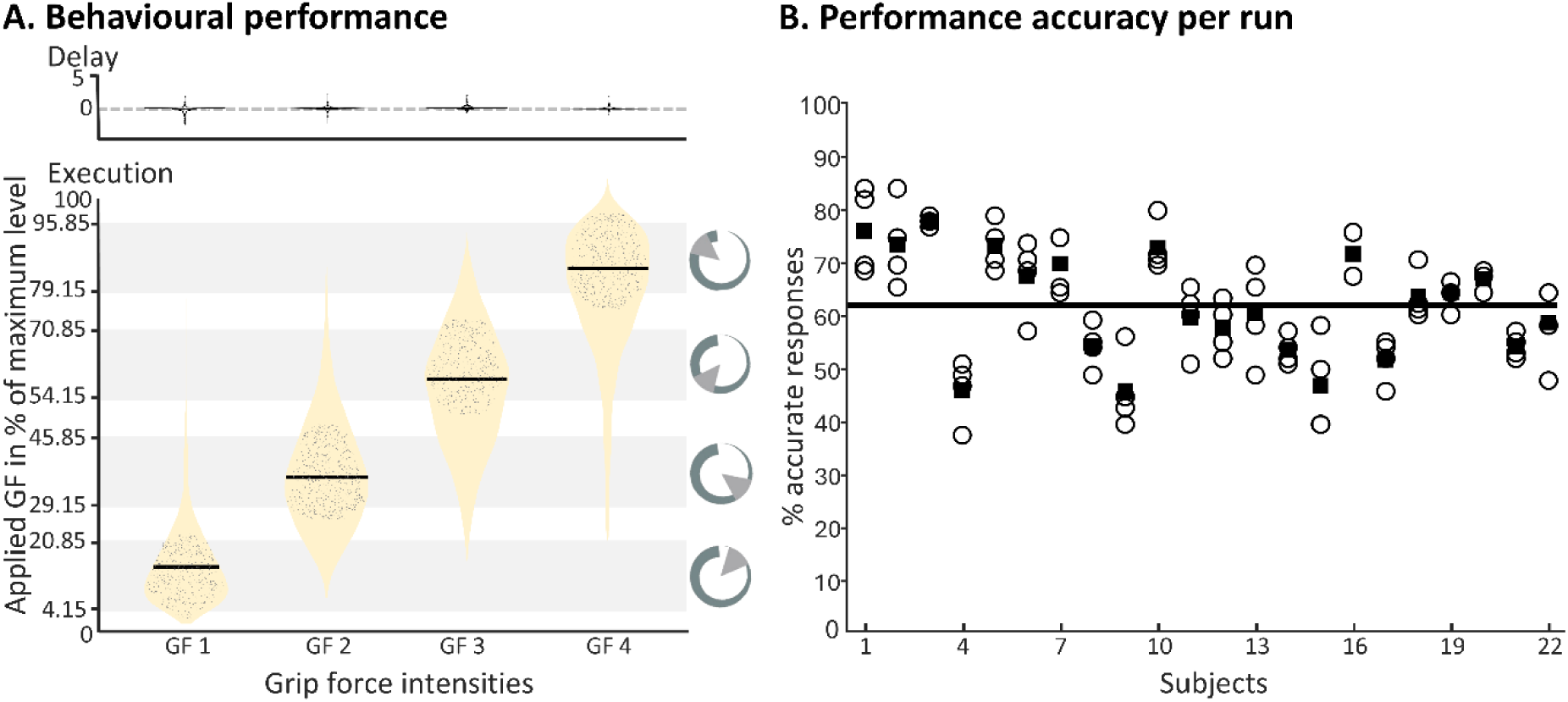
Behavioural assessment. **A.** Displays the grip-forces applied during the delay period (upper display) and the execution period (lower displays). We made sure that only trials were included in the analysis, where participants did not press the grip-force device during the delay periods (upper panel), and grip-force was considered accurate when it did not exceed or fall short by more than 4.15% of the target intensity (lower panel). Light grey bars in the background represent the cued grip-force (GF) levels. For each of the four cued GF levels a violin plot (beige) illustrates the distributions of applied grip-forces, with accurate trial performances indicated as grey dots. **B.** Shows that participants performed broadly consistently across four runs of the delayed grip-force task. Circles represent the performance of accurate grip-force application in each of the four runs (for each participant); filled squares represent the mean performance across runs. The group’s overall mean performance is represented by the black line.

### 2.4 Experimental task

During fMRI scanning, participants performed a delayed grip-force task. Each trial started with the presentation of a *grip-force cue*, comprising of two to-be-maintained grip-force levels presented in cyan and red (illustrated in **Figure 1A**). A 1 s *Effector cue* (cuing the right hand with a right pointer, and the left hand) was presented at 6 s from the beginning of a 12 s *delay period*. The colour of each pointer (either red or cyan) indicated which hand had to be prepared for executing a given grip-force level. The combinations of displayed grip-force levels and the colour of the hand-cues were balanced so that all combinations were presented equally often, and each hand was prepared for each level the same number of times. Upon the display of a *Go-cue*, participants applied simultaneously the anticipated grip-forces with the cued hands; importantly, in the absence of a grip-force or *Effector cue*. A *pre-Go-cue* was presented for 0.5 s before the response to allow optimal response timing during the 1.5 s response window. Finally, visual *Feedback* was provided by showing on a screen the applied forces as green radians if accurate or as red radians if inaccurate, *i.e.* outside of the cued grip-force level (compare **Figure 1A**). The mean grip-force during the last 0.75 s of the response window was evaluated to assess accuracy of responses. For the analysis, trials were considered accurate when the applied force was within the target range ± 15° (4.15% of maximum grip force).

Four experimental runs comprised of 48 experimental trials each. Trial order was fully randomized within a run, where each experimental condition (*i.e.*, the cueing of each of the four to-be anticipated grip-forces with a target hand) was presented equally often (*i.e.*, 12 times per run).

### 2.5 fMRI data acquisition and preprocessing

fMRI data were acquired in 4 runs of 17 min and 2 s each on a 3T Siemens Prisma (Siemens Healthcare GmbH, Erlangen, Germany) at the *Center for Cognitive Neuroscience Berlin* (CCNB) of the *Freie Universität Berlin*. For each run, 1000 functional images were obtained with an EPI sequence (64 channel head coil, 48 slices, interleaved order; TR= 1 sec, 2 × 2 × 2 mm voxel size; multiband acquisition with acceleration factor of 3).

Trial onsets were time-locked to the functional image acquisition to allow a time-resolved analysis (see below). MRI data processing was performed using SPM12, r7771 (*Wellcome Trust Centre for Neuroimaging*, *Institute for Neurology*, *University College London*). To preserve the spatiotemporal structure of the fMRI, data preprocessing was limited to spatial realignment.

#### 2.5.1 Time-resolved searchlight decoding of abstract force intensities and effector-specific grip-force anticipation

We employed an MVPA searchlight approach to identify brain regions that exhibited multivariate parametric codes of force intensity before and after specification of the hands to-be-used. A finite impulse response (FIR) model was used to obtain run-wise beta estimates for each 1 s time-bin of the trials. 20 consecutive time-bins were modelled (as illustrated in **Figure 1C**), comprising 6 time-bins throughout each delay period, 6 time-bins from the *PreGo* cue (i.e., execution period), and 2 time-bins of the ITI. High-pass filtered data (cut-off 192 s) were included in the first-level general linear model (GLM), modelling each combination of intensity and hand. Additionally, six movement parameters, the first five principal components (explaining variance in the white matter and cerebrospinal fluid signals; Behzadi et al., 2007) and a run constant, were added as nuisance regressors, for a total of 688 regressors (8 conditions × 20 time-bins × 4 runs complemented with the 6 realignment parameters and 5 principal components).

To identify where and when information about the intended intensity was encoded in the brain, we applied time-resolved MVPA using SVR (Kahnt *et al*., 2011), which can be considered a multivariate pendant of parametric coding, a method previously applied in a series of WM decoding experiments (*e.g.*, Christophel *et al*., 2012; Schmidt *et al*., 2017; Uluç *et al*., 2020 Pennock *et al*., 2021). Using a searchlight approach (r=4 voxel) independently for every time-bin allows testing for local multivariate representations (*i.e.*, activation patterns of voxels) that code parametric grip-force anticipations. Two SVR models were trained to predict grip-force (*i.e.*, the four grip-force intensities) based on a multivariate data vector (*i.e.*, multivoxel activation pattern) for each hand. To account for differences in the difficulty of grip-force performance, the distances between the four SVR labels were adjusted using Fechner’s law (Fechner, 1860) as previously done (see Caccialupi et al., 2025a), resulting in labels of -2.40, -1.48, 0.45, and 3.42.

Beta estimates of each condition were first normalized (*i.e.*, z-scaled) across the samples for each voxel as implemented in the Decoding Toolbox (TDT; Hebart *et al*., 2015) and forwarded to a four-fold leave-one-run-out cross validation schema. To make our data comparable to previous reports, we used as measure the prediction accuracy defined as the Fisher’s z-transformed correlation coefficient between the predicted value levels and the actual value levels of the test dataset (in TDT: “zcorr”, see also Kahnt *et al*., 2011). The centre of the searchlight was moved voxel wise through the brain, and prediction accuracy values were saved as corresponding whole-brain accuracy maps. In this way, we obtained an accuracy map for every subject and time-bin, reflecting local activation patterns that code abstract or effector-specific grip-force intensity in a multivariate way.

Prediction accuracy maps were entered into a second-level analysis to test for above-chance decoding in terms of SPM’s flexible factorial design implementation of an ANOVA. We used a t-contrast to test for above chance decoding across different time-bins. All results are reported at a threshold of *p* < 0.05 family-wise error (FWE) corrected at the voxel level.

#### 2.5.2 Cross-decoding analysis: Cross-Hand cross-decoding

We conducted a cross-decoding analysis, i.e. cross-hand cross-decoding (*X-Dec: Cross Hand*) to test whether and where parametric representations of grip-force intensity can be found in an effector-independent format (i.e., information about the force intensity is retained in similar brain activation patterns for the two hands). The cross-decoding analysis was based on the same FIR model as the main decoding analysis. The logic underlying the use of the SVR in the cross-decoding, including the four-fold leave-one-run-out cross-validation schema, is the same as in the main analysis, with the only difference being that, in *X-Dec: Cross Hand*, training and testing of the SVR-model were conducted on beta estimates corresponding to different hands. The SVR model was first trained on beta estimates reflecting preparation with the right hand and tested on the left hand, then vice versa (see Caccialupi et al., 2025b).

After normalization and smoothing, prediction accuracy maps were entered into a second-level analysis to test for above-chance decoding using SPM’s flexible factorial design implementation of an ANOVA. This second level design included 1 factor and 20 levels for time-bins (t1–t20). T-contrasts were used to assess above-chance decoding on maps of the second half of each period, as done in the main analysis (accounting for temporal correlation between periods).

An additional second-level analysis was conducted to identify brain regions representing force intensities in similar, effector-independent codes across the first and second delays. For this analysis, we included prediction accuracy maps from the main decoding analyses (i.e., one analysis for hand), which reflect hand-unspecific decoding prior to the *Effector cue* (i.e., during the first delay), as well as from the *X-Dec: Cross-Hand* (covering the first and second delays). The second-level design comprised 3 factors (i.e., one for the right hand, one for the left hand, and one for the cross-decoding between them) and 20 levels corresponding to the time-bins (t1–t20). T-contrasts were computed to assess above-chance decoding on maps reflecting the first delay period (main analyses and cross-decoding), and the second delay (*X-Dec: Cross-Hand*). A conjunction analysis over t-contrasts was performed under the conjunction null hypothesis (Nichols et al., 2005) to identify regions that similarly represent effector-independent information across delays. All results are reported at *p* < 0.05, FWE-corrected at the voxel level.

#### 2.5.3 Test for Temporal Generalization: Cross-Regression Decoding

To test whether multi-voxel activation patterns observed for a specific hand during the second delay period were similar to those during the execution phase, we conducted a cross-regression decoding analysis as a test of temporal generalization. This approach allowed us to assess where in the brain the activation code found during movement execution was represented in a format already present during the preceding preparation phase (see Caccialupi et al., 2025a). In line with previous MVPA studies (e.g., Hebart et al., 2018; Caccialupi et al., 2025a), we tested for temporal generalization by training an SVR on each time-bin statistically assessed at the second level of the corresponding main decoding analysis and testing it on all other time-bins of this set. The cross-regression analysis was based on beta estimates derived from the same FIR model used in the main decoding. To account for temporal correlations of beta estimates across trial periods, SVR training and testing were restricted to the second half of each period (t5-t7, t11-t13, t17-t19 × t5-t7, t11-t13, t17-t19). Two SVRs (one per hand) were trained and tested on these time-bins using a four-fold leave-one-run-out cross-validation scheme, producing 81 cross-regression accuracy maps for the right hand and 81 distinct maps for the left hand.

For the statistical assessment, we averaged prediction accuracy maps to fall into 9 cross-regression accuracy maps (for each hand) to reflect pattern similarities within each combination of time periods. After normalization and smoothing, the cross-regression prediction accuracy maps were entered into two second-level analyses (one for hand) to test for above-chance decoding using SPM’s flexible factorial design specification of an ANOVA. Each design included a first factor with 3 levels for the training periods, and a second factor with 3 levels for the testing periods. We used t-contrasts to assess above-chance cross-regression decoding on combination of accuracy maps reflecting training and testing on each pair of periods, e.g., a single t-contrast was computed over maps reflecting training on the second delay period and testing on the execution period, and vice versa. All results are reported at *p* < 0.05, FWE-corrected at the voxel level.

#### 2.5.4 Control analyses: label permutation control analysis

To control whether prediction accuracies of the main analysis were based on the parametric coding of the grip-force intensity, we performed a label-permutation test. Here, the labels of the conditions entered into SVR-models were permuted. As previously applied (see Schmidt et al., 2017; Uluc et al., 2020; Caccialupi et al., 2025), higher distance of the labelling from the original order should result in reduced decoding accuracies (getting to zero for the highest distance i.e., completely unordered labelling). For all possible permutations, the distance from the rank order was calculated as the sum of the absolute difference of adjacent ranks (e.g., the linear order of grip-force level 1, 2, 3, 4 has a distance of ranks of sum (|1 – 2| + |2 – 3| + |3 – 4|) = 3 and the permuted labelling 2, 1, 3, 4 corresponds to the sum (|2 – 1| + |1 – 3| + |3 – 4|) = 4, resulting in a difference of 1 from the linear order). Thereby, the permutation analysis congregated permutations into four classes of distances from the linear order. For all permutations of labels, the same SVR whole-brain searchlight analyses as in the main decoding were carried out.

To test for above-chance decoding on the maximally permuted label, prediction accuracy maps were entered into the same second-level analysis used in the main decoding. Additionally, to assess whether prediction accuracy increased parametrically with decreasing distance from the linear order of labels, we computed one second-level flexible factorial designs. This included one factor for hand and label type (R-Ordered, R-Distance 1, R-Distance 2, R-Distance 3, R-Distance 4, L-Ordered, L-Distance 1, L-Distance 2, L-Distance 3, and L-Distance 4) and a second factor for time bins (t1–t20). T-contrasts were used to test for above-chance decoding within each time period. All results are reported at *p* < 0.05, FWE-corrected at the voxel level.

#### 2.5.5 Control analyses: HRF-convolved GLM univariate analysis

In addition to the performed MVPA, we also explored univariate activation differences. Specifically, we tested for parametric increases in brain activity during the two delays, and the execution periods. This analysis was based on a classic HRF-convolved GLM, and on functional images that were normalized and smoothed (using SPM’ s default 8 mm FWHM kernel) before entering the first-level model. On the first level, the following regressors were modelled as HRF-convolved boxcar functions for the given phases of the trials: *Delay 1* (*i.e.*, 5–7 s after cue onset), parametric modulator for *Delay 1*, *Delay 2* (*i.e.*, 11–13 s), parametric modulator for *Delay 2*, motor execution period (*i.e.*, 17–19 s), parametric modulator for *execution*, and the motion parameters, as well as the first five principal components (as used in the FIR), as regressors of no interest. For each subject, 12 contrast images were computed (2 hands × 2 modulator types × 3 periods) by averaging beta estimates across runs.

These contrast images were then entered into a single second-level design and one-sample t-test conducted for each condition. All results were reported at *p* < 0.05, FWE-corrected.

## 3 RESULTS

### 3.1 Behavioural performance

The participants included in the decoding analyses (N = 22) responded with accurate application of grip-force, i.e., they were in the target range ± 15° in 62 ± 0.185% (mean ± SD) of the trials. To test for potential differences in performance across hands and grip-force intensities, we conducted a 2 × 4 repeated-measures ANOVA. This analysis revealed no significant main effect of hand (df = 1, F = 1.8959, *p* < 0.1813, η² = 0.0116), but a significant main effect of force intensity (df = 3, F = 67.441, *p* < 0, η² = 0.739), as well as a significant hand × intensity interaction (df = 3, F = 7.7778, *p* < 0.0001, η² = 0.1065). Post hoc t-tests revealed no significant differences in performance between hands at any force intensity. In contrast, for both hands, trials involving the lowest grip-force intensity were slightly easier than those requiring higher intensities. Bonferroni-corrected *p*-values showed that mean performance accuracy with the right hand was significantly higher for intensity 1 compared to intensity 2 (M_diff_ = 33%a, *p* = 0.0001), 3 (M_diff_ = 36%, *p* < 0.0001), and 4 (M_diff_ = 32%, *p* < 0.0001). Similarly, with the left hand, accuracy for intensity 1 was significantly higher than for intensity 3 (M_diff_ = 20%, *p* = 0.0018) and 4 (M_diff_ = 21%, *p* < 0.0004). These findings are consistent with those of a previous study by Caccialupi et al. (2025a). Accordingly, we applied an adjustment based on Fechner’s Law to model the subjectively perceived differences in grip-force intensity for the neuroimaging analyses (see Caccialupi et al., 2025a). Violin plots in **Figure 2A** display the distribution of responses (*i.e.*, accurate and non-accurate performances) in terms of applied force on the grip-force device for the four grip-force intensities (averaged between hands, due to not significant differences). We carefully ensured that the included participants did not apply force during the delay period (see **Figure 2A**, upper display).

To assess potential training or fatigue effects across the four runs, we conducted a 1 × 4 repeated-measures ANOVA (averaging performance accuracies across hands), which did not reveal differences (df = 3, F = 0.0079, *p* = 0.999, η2 = 0.0001. Performance accuracy per participant and run is illustrated in **Figure 2B**.

### 3.2 Multivariate mapping of regions that code abstract intensity and effector-specific grip-force anticipation

To identify regions parametrically coding force intensities in two delay periods, we conducted a time-resolved whole-brain SVR analysis. Within a second-level ANOVA design, we computed t-contrasts across prediction accuracy maps for the two trial periods of interest: *Delay 1* (t5–t7) and *delay 2* (t11–t13). T-contrasts were computed on the second half of each period to account for temporal correlations in the BOLD signal and were shifted by one time-bin toward *execution* to better account for latency in the BOLD response. All results are reported at *p* < 0.05, FWE-corrected. During *Delay 1* (i.e., before instructing which hand must be prepared for which intensity), we computed a single t-contrast for the two hands and found strong above-chance decoding in the praecuneus (PCu) (see **Figure 3A**, first column, and **Table 1**). During the second delay, two lateralized clusters comprising contralateral intraparietal sulci (pIPS/aIPS), somatosensory cortex (S1), dorsal premotor cortex (PMd), and supplementary motor area (SMA) were revealed in the right and left hemispheres during preparation with the left and right hands, respectively (see **Figure 3A**, second column, and **Table 1**). Finally, a control t-contrast on the motor execution period (i.e., t17-t19) showed expected bilateral primary motor cortices (M1s), with highest significance in the respective contralateral hemispheres (**Figure 3A**, **Table 1**).

**Figure 3.**
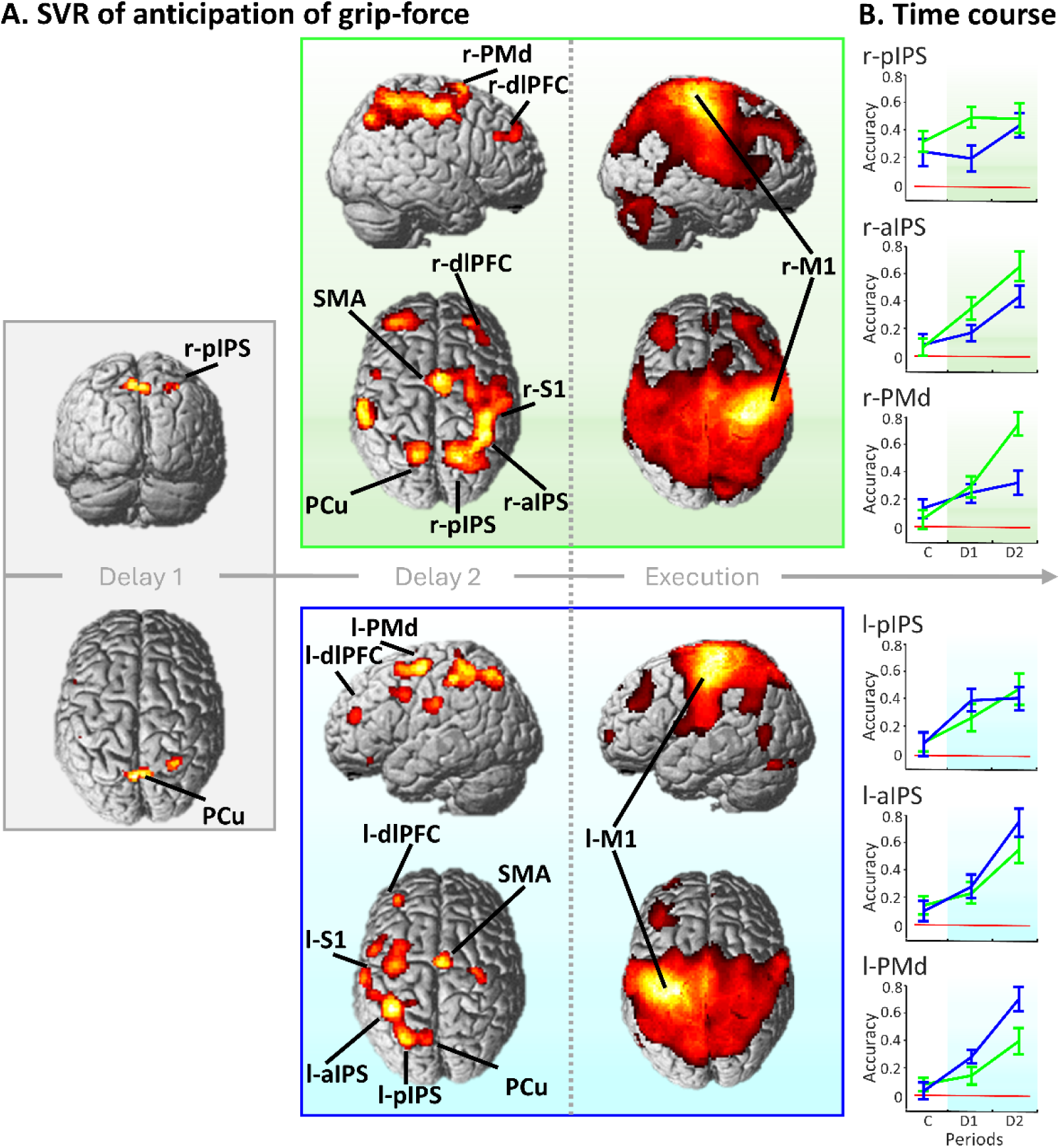
Results of time resolved support vector regression analyses **A.** Displays brain regions that parametrically code grip-force intensities during the first delay period (Delay 1), second delay period (Delay 2) and motor execution period (Execution). The grey panel indicates encoding of force intensities regardless the hand to-be used (Delay 1); the upper, green panel refers to preparation with the left hand (Delay 2); the lower, blue panel, preparation with the right hand (Delay 2). Brain regions with above-chance prediction accuracy are revealed by t-contrasts testing the respective periods against zero, at *p* < 0.05, FWE-corrected. In Delay 1, two clusters were found in the PCu and r-pIPS (first column). In Delay 2 (second column), two lateralized clusters were revealed in the pIPS, aIPS, S1, PMd, and SMA, on contralateral hemispheres respectively (and the addiction of similar ipsilateral patterns in the PCu). Finally, t-contrasts on the Execution revealed contralateral and ipsilateral clusters in the M1s (third column). **B.** Time-courses of prediction accuracy for left- and right-hand preparation (represented in green and blue) were extracted from peak voxels identified in the main analysis (Delay 2), in contralateral r-pIPS [*x*=26, *y*=--62, *z*=50], l-pIPS [*x*=-28, *y*=--66, *z*=52], r-aIPS [*x*=-36, *y*=-42, *z*=58], l-aIPS [*x*=-38, *y*=-42, *z*=58], r-PMd [*x*=30, *y*=-12, *z*=56] and l-PMd [*x*=-38, *y*=--8, *z*=62]. Please note that because time-courses were extracted from the most significant voxels they are only displayed for descriptive purpose. No further statistical testing was conducted to avoid circular conclusions.

**Table 1.** Regions that exhibit above-chance prediction accuracy across the first and second delay periods and motor execution period, revealed by a t-contrast displayed at *p* < 0.05, FWE corrected.

| Cluster size | Anatomical Region | Peak MNI coordinates |  |  | z-score |
| --- | --- | --- | --- | --- | --- |
|  |  | x | y | z |  |
| First delay Period |  |  |  |  |  |
| 201 | PCu | -4 | -66 | 56 | 5.04 |
| 97 | Right pIPS | 28 | -56 | 54 | 4.95 |
| Second delay Period, left hand |  |  |  |  |  |
| 6073 | Right pIPS | 26 | -62 | 50 | 7.33 |
|  | Right aIPS | 36 | -42 | 58 | 6.88 |
|  | SMA | 2 | -2 | 58 | 6.47 |
|  | Right PMd | 30 | -12 | 56 | 6.39 |
| 924 | Left aIPS | -58 | -28 | 34 | 6.87 |
| 860 | Left pIPS | -16 | -60 | 56 | 5.82 |
| 325 | Right dIPFC | 30 | 48 | 32 | 5.68 |
| 648 | Left dIPFC | -34 | 48 | 24 | 5.20 |
| 79 | Left PMd | -50 | 4 | 42 | 4.86 |
| Second delay Period, right hand |  |  |  |  |  |
| 1716 | Left pIPS | -28 | -66 | 52 | 6.32 |
|  | Left aIPS | -38 | -42 | 58 | 5.88 |
| 322 | SMA | 4 | -4 | 66 | 5.89 |
| 688 | Left PMd | -38 | -8 | 62 | 5.77 |
| 150 | Left dIPFC | -34 | 46 | 20 | 5.61 |
| 245 | Left S1 | -62 | -16 | 28 | 5.47 |
| 172 | Right PMd | 34 | -16 | 54 | 5.11 |
| 202 | Left PMv | -54 | 6 | 38 | 5.10 |
| Motor execution Period, left hand |  |  |  |  |  |
| 39062 | Right M1 | 30 | -26 | 62 | 19.57 |
| 583 | Right S1 | 38 | -22 | 52 | 5.59 |
|  | Left M1 | 30 | 28 | 58 | 5.93 |
| Motor execution Period, right hand |  |  |  |  |  |
| 25730 | Left M1 | -32 | -26 | 60 | 16.99 |
|  | Left S1 | -40 | -36 | 54 | 10.44 |
|  | Right M1 | 34 | -36 | 58 | 5.66 |

To illustrate the temporal evolution of decoding accuracy across the three trial periods, we extracted time courses of prediction accuracy from the peak voxels in the pIPS, aIPS, and PMd, as found in the group-level statistics during *Delay 2* (see **Table 1** for MNI coordinates). Each time course represents the mean prediction accuracy for each period, averaged across the respective time-bins (as previously done by Caccialupi et al., 2025b; see **Figure 3B**). For each hand, the second delay period showed a stronger increase in prediction accuracy in the contralateral hemisphere, reflecting parametric coding in effector-specific regions (i.e., areas showing systematic lateralization of above-chance decoding for the to-be used hand). This pattern supports the high specificity of the observed effects and indicates effector-specific coding of anticipated grip-force intensities in contralateral brain-regions.

Together, these results likely identify regions where information is parametrically represented in an effector-specific format, following an earlier, abstract representation of force intensities in the PCu. Each of the reported regions showed no difference in prediction accuracy between hands during the first delay period. All regions, except for the PCu, exhibited a substantial increase in decoding accuracy during motor execution.

### 3.3 Testing for abstract, effector-independent representation by cross-decoding

To test where in the brain parametric codes of grip-force intensity are represented in an abstract, effector-independent format, we conducted a whole-brain cross-decoding analysis. Force intensities were decoded using an SVR-model trained on beta estimates from one hand and tested on the other, and vice versa. In a second-level ANOVA design, t-contrasts were computed for each delay on maps corresponding to the same time bins tested in the main analysis. Consistent with the main decoding results, we identified a single cluster spanning the bilateral PCu, showing robust above-chance decoding within each delay (at *p* < 0.05, FWE-corrected; see **Figure 4A** and **Table 2**). While above-chance decoding during the first delay likely reflects abstract, effector-independent coding of force intensity (as also indicated by results of additional SVR decoding, and cross-decoding analyses; see **Figure S1** in the *Supplementary Materials)*, it could alternatively indicate simultaneous coding of information about both hands during the second delay.

**Figure 4.**
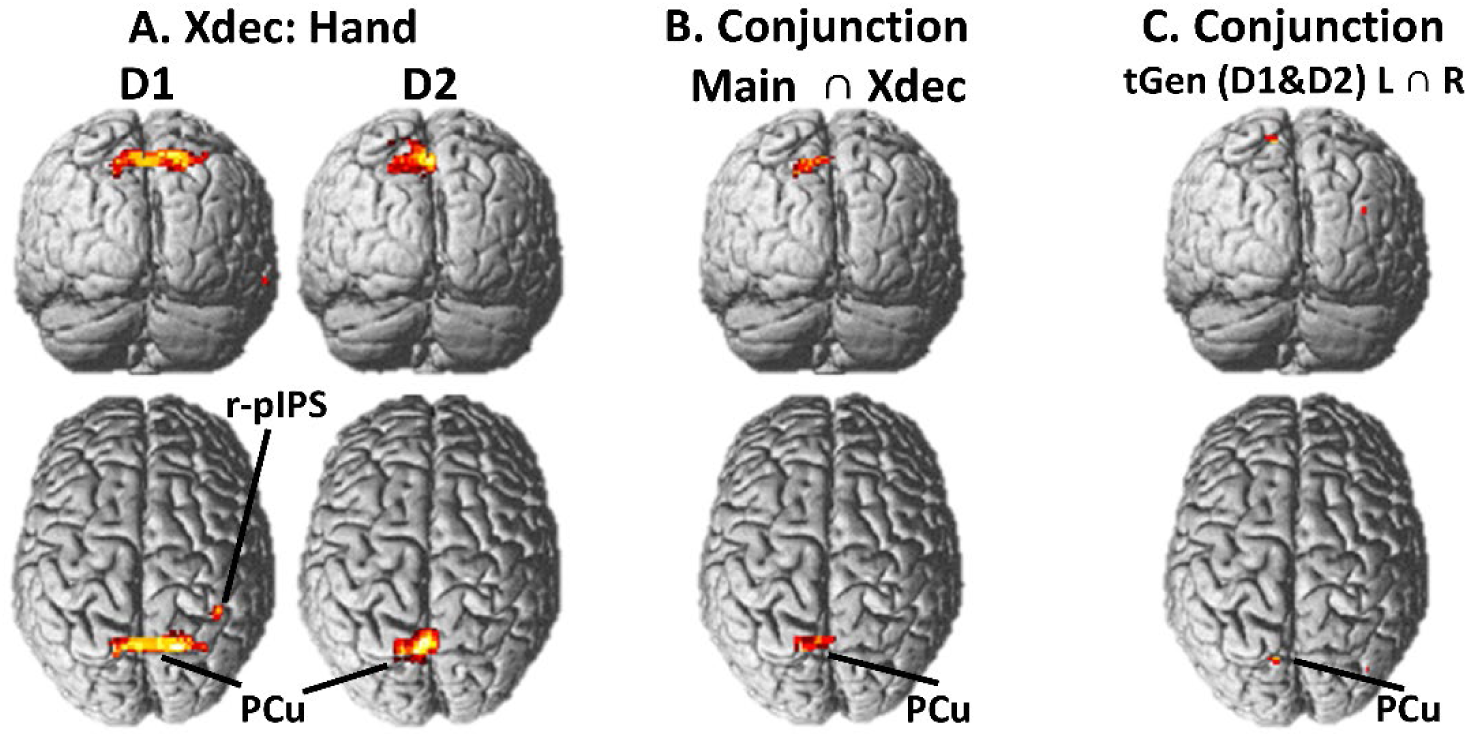
Cross-decoding analyses testing abstract format representation of force intensity **A.** Displays results of cross-effector decoding (Xdec: Hand) for Delay 1 (D1; first column) and Delay 2 (D2; second column). Brain regions with above-chance prediction accuracy are revealed by t-contrasts testing the respective periods against zero, at *p* < 0.05, FWE-corrected. During both delays, we found a similar cluster in the PCu, as also found in the main decoding (D1). **B.** Conjunction analysis over three t-contrasts (Xdec: Hand D1, Xdec: Hand D2, and the main decoding D1) (Conjunction Main ∩ Xdec) was conducted under the conjunction null hypothesis (at *p* < 0.05, FWE-corrected) to test whether an abstract format representation of force intensity is present during the two delay periods. In both delays, the analysis revealed a significant activation pattern in the PCu, similar to those found in Xdec: Hand (D1 and D2) and in the main decoding (D1). **C.** An additional conjunction analysis over t-contrasts was conducted to test for temporal generalization between D1 and D2, i.e., Conjunction tGen (D1&D2) L∩R. This analysis was based on accuracy maps derived from cross-regression decoding performed separately for the two hands. The conjunction of t-contrasts for the two hands (L∩R), revealed a small pattern in the PCu, which is included in all the analyses testing effector-independent coding of force intensities. Together with Xdec: Hand D2, this finding indicate that an abstract format representation of grip-force intensity is maintained in the PCu across the two delay periods.

**Table 2.** Regions that exhibit above-chance prediction accuracy across the first and the second delay periods (Xdec: Hand), and between them (as revealed by conjunction with the main decoding, and temporal generalization; tGen), at *p* < 0.05, FWE corrected.

| Cluster size | Anatomical Region | Peak MNI coordinates |  |  | z-score |
| --- | --- | --- | --- | --- | --- |
|  |  | x | y | z |  |
| ***Xdec: Hand, first delay Period*** |  |  |  |  |  |
| 486 | PCu | 0 | -62 | 54 | 5.31 |
| 28 | Right pIPS | 40 | -44 | 54 | 4.80 |
| ***Xdec: Hand, second delay Period*** |  |  |  |  |  |
| 477 | PCu | -2 | -60 | 54 | 5.52 |
| ***Conjunction: Main, D1 ∩ Xdec: Hand, D1 ∩ Xdec: Hand, D2*** |  |  |  |  |  |
| 170 | PCu | -2 | -62 | 56 | 5.23 |
| ***Conjunction: tGen D1&D2, L ∩ tGen D1&D2, R*** |  |  |  |  |  |
| 7 | PCu | -8 | -72 | 64 | 4.59 |

To further test whether the above-chance decoding in *Delay 2* reflects abstract codes of force-intensity, we cojoined results of the *X-Dec: Cross Hand* analysis from *Delay 2* with those from *Delay 1*, and the main decoding analysis (first delay). Within a second-level ANOVA design, t-contrasts were computed on prediction accuracy maps for the same time periods tested in the main decoding. The conjunction analysis (tested against a conjunction null hypothesis; Nichols et al., 2005) revealed a similar pattern in the PCu as compared to those found in the cross-decoding (within *Delay 1*), at *p* < 0.05 FWE-corrected (**Figure 4B**). This result suggests that parametric codes in the PCu represent force intensities in a similar format across both delay periods. Given that the intensities are initially encoded in an abstract, effector-independent format within the PCu, it is thereby likely that a similar code during the second delay represents force-intensities regardless the hand to-be-used.

Taken together, the *X-Dec: Cross Hand* and the main decoding analyses provide converging evidence for the abstract, namely effector-independent, format representation of parametric information about force-intensities in the PCu.

### 3.4 Test for temporal generalization

To assess whether multivariate patterns of grip-force intensities observed during *Delay 1* and *Delay 2* were similar to those during motor execution, we performed whole-brain cross-regression decoding. All combinations of time periods were tested by first training on early time bins and testing on later time bins, and then vice versa. Within a second-level ANOVA design, t-contrasts were independently computed for each hand on pairs of cross-regression accuracy maps, with the first map reflecting training on the earlier period and the second on the later. All results are displayed at *p* < 0.05, FWE-corrected, with green background indicating the left hand and blue background indicating the right (see **Figure S1** in the *Supplementary Materials*).

First, to test for generalization between *Delay 1* and *Execution*, we computed two independent t-contrasts for the left and right hands. These contrasts revealed no brain regions with above-chance decoding, even at an uncorrected threshold of *p* < 0.001 (see **Figure S2**, first column, and **Table S2**). This suggests that the initial abstract representation of force intensities may be transformed into a different code during motor execution. Second, cross-regression decoding was performed between the first and second delays, revealing above-chance decoding in the PCu for both the left and right hands (see **Figure S2**, second column). Notably, an additional conjunction analysis (tested against the conjunction null hypothesis) identified a small cluster in the PCu, included in all the above-mentioned analyses (see **Figure 4C** and **Table 2**). Together, these findings indicate that an abstract format representation of grip-force intensity is maintained across the two delay periods. Finally, when testing for generalization between *Delay 2* and *Execution*, we found two lateralized patterns in the IPS and PMd, located in the right and left hemispheres for the left and right hands, respectively (see **Figure S1**, third column). This indicates that, over time, the neural representation of intended force intensity may shift from an abstract format representation (in the PCu) to a movement-specific code involving contralateral IPS and PMd, i.e., regions exhibiting similar activation patterns during preparation and execution.

### 3.5 Control analyses: Label permutation test

To corroborate specificity of the main analysis, we conducted label permutation testing and two second-level analyses. First, we tested above-chance decoding for the maximally permuted labels in terms of SPM’s flexible factorial design implementation of an ANOVA. When t-contrasts are computed on the first and second delay periods (as in the main analysis), we found no significant clusters throughout the whole brain with identical *p* < 0.05, FWE correction. The same result was observed when inspected at *p* < 0.001 uncorrected. For illustrative purposes, we display the time-courses of the label-permutation tests with increasing dissimilarity to the original order for the peak voxels of the main analysis (see **Figure S3B** in the *Supplementary Materials*). As expected, the time-course of the completely unordered labelling do not show above chance prediction-accuracies throughout all phases of the experimental trials.

As a second control analysis, we entered prediction accuracy maps resulting from permutation labelling into a second-level flexible factorial designs (as in the main analysis). A single second level design included one factor for hand and label types (i.e., R-Ordered, R-Distance 1, R-Distance 2, R-Distance 3, R-Distance 4, L-Ordered, L-Distance 1, L-Distance 2, L-Distance 3, and L-Distance 4), and a second factor for time bins (t1–t20). We computed parametric contrasts over maps corresponding to the same time bins tested in the main analysis to assess whether prediction accuracies in similar brain regions showed a parametric decrease with increasing dissimilarity of labels from the original intensity order. When assessed at *p* < 0.05, FWE-corrected, the results corroborated the main findings, showing highly similar clusters across all three time periods (see Figure **S3A** and **Table S3** in the *Supplementary Materials*). This indicates that the above-chance prediction accuracies observed in the main analysis were likely driven by the parametric coding of grip-force intensity.

### 3.6 Control analyses: HRF convolved GLM

To test for parametrically modulated activation strength during the two delay periods and motor execution, we performed a univariate analysis based on an HRF convolved first level GLM model. One sample t-tests were computed to assess group level effects of first-level contrasts, showing no significant parametric modulation during the first and second delays (at *p* < 0.05, FWE-corrected). During *Execution*, only the contralateral r-M1 and l-M1 exhibited parametric activity modulation (see **Figure S4A**, and **Table S4** in the *Supplementary Materials*). These results make it rather unlikely that parametric effects found in the main MVPA analyses were primarily driven by univariate effects.

## 4 DISCUSSION

In this fMRI study, we used a delayed grip-force task in combination with time-resolved SVR to identify brain regions that parametrically code grip-force intensities. Specifically, we tested whether, during a first delay period (i.e., prior to effector specification), brain activity patterns represent intended force intensities in an abstract, effector-independent format, and how these representations are subsequently transformed into effector- and movement-specific codes during a second delay.

To this end, we adapted our previous delayed grip-force paradigm (Caccialupi et al., 2025a, 2025b) to include two consecutive delay periods: during the first, participants maintained two of four possible force intensities for later execution; during the second, they were cued which hand to prepare for each force. This design allowed us to test the abstract, effector-independent coding of force intensity during the first delay and its subsequent transformation into effector-specific representations during the second. During the first delay (lasting 6 s from trial onset), graded grip-force intensities could be decoded from activity patterns in the PCu. Meanwhile, during the second delay (lasting 6 s from effector-cue onset), above-chance decoding was found in the contralateral aIPS and PMd, localized to the right hemisphere for the left hand and to the left hemisphere for the right. This systematic lateralization indicates that anticipated grip-force intensities are represented in effector-specific brain regions during the second delay. Notably, the main decoding analysis also revealed that graded grip-force intensities are parametrically represented in distinct areas across the two delays. Together, these findings suggest that delay-specific activation patterns may reflect successive stages of motor planning: the encoding of a motor goal (i.e., the intended force intensity) in an abstract format during the first delay and its subsequent transformation into effector-specific movement plans during the second.

To further characterize the change in representational formats across delays, we applied cross-decoding and generalization tests to determine when and where force intensities were represented in stable representational codes (i.e., same activity patterns which allow above-chance cross-decoding or generalization) and where activation patterns are only exhibited in a condition-specific or time-specific manner. First, we performed cross-effector decoding (*Xdec: Hand*). Here, the SVR model was trained and tested on different hands. This analysis revealed above-chance decoding in the PCu during both delays. Coherently, a conjunction analysis of t-contrasts (*Xdec: Hand* in D1 and D2, and main decoding in D1) identified overlapping clusters in the PCu, confirming that this region encodes abstract features of intended force intensities, even into the second delay. Consistently, cross-regression decoding indicated that parametric codes in the PCu remained stable across delays, whereas contralateral aIPS and PMd exhibited stable activation patterns from the second delay through execution. Thereby, parametric codes in the aIPS and PMd are likely represented in a movement-specific format that emerges from the second delay onward. Together, these results indicate a progressive transformation in representational format, from abstract coding in the PCu to effector-and movement-specific coding in aIPS and PMd.

These results align well with the two-stage framework of action planning (Boettcher et al., 2021), which proposes that, following the initial encoding of a motor goal, the intended action is specified into effector-specific movement plans. Accordingly, in our study, the PCu likely encoded grip-force intensities (i.e., the motor goal) in an abstract format, contributing to the early stage of motor planning (Hummel et al., 2003; Astafiev et al., 2004). As planning progressed, contralateral aIPS and PMd were likely engaged in effector selection and in anticipating the movements required to apply the intended intensity, even before execution begins (Gale et al., 2021; Diedrichsen et al., 2022). Within PMd, specific movement parameters, such as the muscle fibres recruited for later grip-force execution, may have been represented and refined in preparation for movement onset (Mizuguchi et al., 2013; Caccialupi et al., 2025a, 2025b).

In the following sections, we discuss our findings in temporal sequence, from the first to the second delay period.

### 4.1 The First Delay Period: Action Specific encoding of Graded Force Intensities in the Precuneus, and Modality-Independent Representation in the Right Intraparietal Sulcus

During the first delay period, our main decoding analyses revealed two significant clusters, one in the PCu and the other in the r-pIPS.

The most pronounced cluster was located in the PCu, a region well known for its role in planning motor actions (Gallivan et al., 2011, 2015; Wang et al., 2019). Previous time-resolved MVPA studies using delayed-response paradigms have coherently shown that the PCu is involved in planning hand movements during the delay period preceding the execution of goal-directed actions (Soon et al., 2008; Bode & Haynes, 2009; Gallivan et al., 2011). Crucially, above-chance decoding in the PCu has also been found during an early phase of this delay, preceding effector specification (Soon et al., 2013). While this finding has been interpreted as evidence that the PCu encodes abstract intentions independent of effector-specific plans (Soon et al., 2013), such abstraction does not imply a functional detachment from motor control (Gallivan et al., 2011). Rather, early increases in prediction accuracy in the PCu, together with task-dependent activation patterns in delayed-response paradigms (when abstract information is maintained and transformed for later execution of complex motor actions) more frequently reflect higher-order representations of action features within the motor hierarchy (Hamilton & Grafton, 2007; Gertz et al., 2017; Ariani et al., 2018; Monaco et al., 2020).

Consistent with these results, recent fMRI MVPA studies have revealed a medial-to-lateral gradient of abstraction within the PPC (Heed et al., 2013, 2016). Accordingly, medial regions such as the PCu have been found to encode effector-independent aspects of motor plans (e.g., motor goals; Heed et al., 2013), whereas more lateral regions, including the aIPS, encode effector-specific representations of the intended action (Heed et al., 2016; Caccialupi et al., 2025b). Moreover, converging evidence indicates that a dorsomedial parieto-frontal circuit linking the PCu with the aIPS and PMd is specialized for grasp planning (Heed et al., 2013; Gallivan & Culham, 2015), and exhibits a posterior-to-frontal gradient of abstraction. Within this circuit, posterior regions such as the PCu were found to represent the intended action (i.e., the motor goal) regardless of movement details (Errante et al., 2021), whereas the PMd encodes detailed motor parameters in preparation for execution (Caccialupi et al., 2025a). Furthermore, functional coupling between the PCu and aIPS emerges when effector selection is required (Wang et al., 2019), and above-chance decoding in the contralateral aIPS is observed as the hand used to apply a grip-force intensity is specified (Caccialupi et al., 2025b). Taken together, these results indicate that PPC subregions differentially code abstract and effector-specific information during an early planning phase of grasping actions (Gallivan & Culham, 2015; Caccialupi et al., 2025b). Coherently, our findings further show that parametric grip-force intensities can be decoded from the PCu during the first delay: in an abstract, effector-independent format, preceding effector- and movement-specific planning in the contralateral aIPS and PMd.

Besides the PCu, during the first delay, we also found above-chance decoding in the r-pIPS. This region has been repeatedly implicated in representing abstract quantities including numerical magnitude, size, and other continuous variables (Dehaene et al., 2003; Piazza et al., 2004; Castelli et al., 2006; Eger et al., 2009). Recent fMRI MVPA studies also suggest that parametric modulation in the r-pIPS reflects the encoding of graded abstract quantities (Pennock et al., 2021). Unlike the PCu, whose parametric codes likely represents abstract action features, the r-pIPS may provide a modality-independent representation of abstract parameters that can be flexibly accessed for both cognitive and motor functions. Accordingly, during the first delay, parametric codes in the PCu and r-pIPS may respectively reflect modality-specific (action-related) and modality-independent neural representations of graded intensities, highlighting the simultaneous processing of motor-relevant information across PPC subregions.

### 4.2 The Second Delay Period: Effector Selection in the Anterior Intraparietal Sulcus and Movement Planning in the Dorsal Premotor Cortex

During the second delay period, SVR decoding revealed significant, lateralized clusters in the contralateral IPS and PMd; two regions well established for their roles in motor planning.

The most prominent clusters were located in contralateral sections of the aIPS, with parametric codes specific for force intensities to-be prepared with the right hand in the l-aIPS, and codes specific for the left hand in the r-aIPS. Consistent with this, previous fMRI MVPA studies have shown that during grasp planning, the contralateral aIPS predominantly encodes information specific to a single effector, i.e., the hand to be used (Heed et al., 2016). Moreover, in delayed grip-force tasks, above-chance decoding systematically lateralizes to the hemisphere contralateral to the prepared hand and reflects action-specific properties such as grip-force intensity (Caccialupi et al., 2025b). This contralateral lateralization, which emerges in regions specific to the body part being used, is considered a reliable indicator that the aIPS represents effector-specific information for hand actions (Gallivan et al., 2011). While abstract or movement-specific features of planned actions can also be decoded from the pIPS regardless of effector, the aIPS primarily supports effector-specific representations during motor planning (Gallivan et al., 2013). Consistent with this, MVPA and SVR analyses have shown that detailed movement parameters—such as grip type and force intensity—are encoded in these effector-specific regions (Ariani et al., 2015; Caccialupi et al., 2025). The effector sensitivity of the aIPS has been further demonstrated across effectors: successful decoding of movement direction occurs for hand movements but not for saccades, and cross-decoding between them did not yield above-chance accuracy (Gallivan et al., 2013). This functional specialization, together with systematic lateralization, indicates that the contralateral aIPS contributes to representing movement plans in an effector-specific code (Gallivan & Culham, 2015).

Coherently, time-resolved decoding analyses have revealed faster increases in prediction accuracy for hand actions in the pIPS and PCu compared to the contralateral aIPS and PMd (Gallivan et al., 2011; Ariani et al., 2015), implicating the aIPS in a later stage of motor planning—such as effector selection and movement specification (Caccialupi et al., 2025b). A similar temporal progression has been observed when motor planning involves grip-force anticipation (Caccialupi et al., 2025a). In this context, the initial contribution of the pIPS in encoding parametric codes of grip-force intensities has been attributed to a transformation process—from abstract representations of intended force intensities into concrete movement plans encoded in the PMd. Given the extensive anatomical and functional connections among the PCu, pIPS, and aIPS that links these regions to the PMd in a dorsomedial parieto-frontal circuit involved during grasp planning (Monaco et al., 2021), the aIPS may serve as an intermediate cortical hub that transforms abstract information about grip force into effector-specific codes. Receiving input from the pIPS and PCu, the aIPS likely contributes to specifying motor goals into detailed movement plans by representing the intended action in an effector-specific format (Leoné et al., 2014; Caccialupi et al., 2025b). In line with this, our results indicate that during the second delay, the aIPS parametrically represents anticipated grip-force intensities in an effector-specific (body-centered) reference frame, which is essential for subsequent movement planning in the PMd.

Robust decoding was also found in the contralateral PMd, a region well known for its contribution to movement planning (Cisek & Kalaska, 2005; Gallivan et al., 2016). In fMRI MVPA studies, the PMd is most frequently found to encode information specific to planned movement features (Gallivan & Culham, 2015), e.g. the grip force and the grip-type (Ariani et al., 2015; Caccialupi et al., 2025a). Coherently, within the fronto-parietal network involved in motor planning, the PMd is regarded as a downstream node mediating the transfer of action-related information and movement plans to M1. In particular, psychophysiological interaction (PPI) analyses of fMRI data have shown that during a delay period preceding execution, the PMd receives input from parietal regions—most notably the aIPS—and transmits signals to M1 (and possibly S1) in a movement-specific, task-dependent manner (Monaco et al., 2021). During motor planning, PMd activation patterns are most frequently found to represent movement-specific information in body-centered coordinates rather than in an extrinsic reference frame (Klautke et al., 2023). In virtue of this format representation, the PMd is believed to exert control over the M1, and corresponding muscle groups through top-down corticospinal signalling (Mizuguchi et al., 2011). Notably, combined fMRI–EMG studies support this interpretation: during motor imagery of graded grip-force application, PMd activity and muscle activation exhibit similar parametric modulation (Mizuguchi et al., 2011, 2013), suggesting that detailed motor components—such as the desired state of muscle groups—can be specified at higher cortical levels (Gordon et al., 2023). To clarify the PMd’s role in representing the intensity of upcoming grip force during motor planning, van Nuenen et al. (2012) combined repetitive transcranial magnetic stimulation (rTMS) with fMRI in a delayed grip-force task. Their findings indicate that rTMS over the PMd—but not M1—modulates the influence of predictive cues on grip-force execution, highlighting the PMd’s critical role in anticipating graded grip-force intensities. Compared to M1, which is primarily engaged during movement execution, the PMd is most likely involved in representing detailed motor parameters, such as the intended force intensity, during movement planning (van Nuenen et al., 2012; Caccialupi et al., 2025a).

Coherently, time-resolved SVR and cross-decoding analyses reveal that parametric codes in the PMd (specific to anticipated grip-force intensities) are represented in brain activation patterns that during a late delay phase become increasingly similar to movement-specific patterns found during execution (Caccialupi et al., 2025a). As even found in the current study, this result indicates that movement specific information might be encoded in a movement-specific format before execution. At last, fMRI MVPA studies show that the contralateral PMd encodes grip types in an effector-specific format (Ariani et al., 2018). Together, these findings suggest that a key function of the PMd might rests in the integration and further specification of effector-specific information (initially encoded in the aIPS) into detailed movement plans, anticipating concrete motor parameters—such as muscle contraction levels—required for forthcoming execution.

Taken together, our results indicates that, after the initial encoding of more abstract features of the intended force intensity (in the PCu), parametric codes in the aIPS and PMd (found during the second delay) may carry information specific for the effector and the concrete motor parameters to-be controlled during a forthcoming motor execution phase, i.e., the anticipated grip-force intensity.

### 4.3 Cross-Decoding Analyses: from Abstract to Effector- and Movement-Specific Force Intensity coding

To investigate how intended force intensities were represented in the brain, and how representational formats changed across delay periods, we performed two complementary cross-decoding analyses. The first, cross-effector decoding (*Xdec: Hand*), assessed whether (in the brain) parametric activity patterns encoded force intensities in an abstract, effector-independent format, with first and second delay examined separately. The second, cross-regression decoding (used to test for temporal generalization), evaluated the stability of these activity patterns, including their similarity between the execution period and earlier delays.

The *Xdec: Hand* was conducted using a searchlight approach and a time-resolved SVR-model, trained on beta estimates from one hand and tested on the other. Consistent with the main decoding results, it revealed a robust significant cluster in the PCu during both delay periods (at *p* < 0.05, FWE-corrected). During the first delay, above-chance decoding likely reflected abstract, effector-independent coding of force intensity (as no information about the effector was available in this period), whereas during the second delay, similar activation patterns for the two hands may indicate a shared representation (with information about each hand stored in similar activation patterns). To further determine whether force intensities were represented in an abstract format during the second delay, we performed a conjunction analysis testing the overlap of activation patterns found in the main decoding (*Delay 1*) and cross-effector decoding (both delays) against a conjunction null hypothesis (Nichols et al., 2005). The resulting PCu cluster (tested at *p* < 0.05, FWE-corrected) largely overlapped with those identified during the first delay, confirming that parametric activation patterns in the PCu encode force intensities in a consistent, abstract format across delays. This suggests that, while the second delay shows effector-specific codes in contralateral aIPS and PMd, the PCu represents intended force independently of the selected hand. A small cluster in the right pIPS also showed above-chance cross-hand decoding during the first delay, consistent with its supporting role in abstract quantity representation.

Cross-regression decoding further examined the generalization of parametric codes in the PCu between the first and second delays. Specifically, independent tests were conducted for each hand, followed by a conjunction analysis across hands to evaluate the similarity of abstract force-intensity codes between delays. This procedure revealed a cluster consistent with those identified in the main and cross-effector decoding analyses (at *p* < 0.05, FWE-corrected), indicating that abstract representations of intended force intensities are maintained in the PCu. Moreover, no above-chance decoding was observed between the first delay and the execution period, even when tested at a more liberal threshold of *p* < 0.001, uncorrected. This finding suggests that abstract representations in the PCu are reformatted into a different representational format during execution. In contrast, cross-regression decoding between the second delay and the execution period revealed lateralized patterns in contralateral aIPS and PMd - regions previously identified in the main decoding analysis (at *p* < 0.05, FWE-corrected). Because activation patterns during grip-force execution reflect detailed movement parameters, the presence of similar parametric codes during the second delay suggests that anticipated grip-force intensities are already encoded in a movement-specific format (as also found in Caccialupi et al., 2025a). Taken together, these results indicate a progressive shift in representational format—from an abstract, effector-independent representation in the PCu to effector- and movement-specific codes in the aIPS and PMd that closely resemble movement-related activation patterns.

Overall, cross-decoding analyses provide converging evidence for a two-stage transformation process of force-intensity representations. Specifically, we found that abstract representations of intended force intensities are initially encoded in the PCu, persist across delays, and are ultimately transformed into effector- and movement-specific codes in the aIPS and PMd during the second delay. This temporal progression highlights the distinct yet complementary roles of PPC subregions and the dorsal premotor cortex in transforming abstract representations of force intensities into executable movement plans - that is, anticipated grip-force intensities.

### 4.4 Control analyses

The specificity of our decoding results was further corroborated by three control analyses.

First, we performed supplementary SVRs and cross-decoding analyses to test whether force intensities were encoded in an abstract format during the first delay period, independent of the color in which the force levels were presented in the *grip-force cue* (see *Supplementary Materials*, **Figure S1** and **Table S1**). In the SVRs, beta estimates were entered into two separate SVR-models sorted by level color, allowing us to account for the potential influence of level color on force intensity coding. A t-contrast over prediction accuracy maps for cyan and red presentations revealed a significant cluster in the PCu (*p* < 0.05, FWE-corrected), consistent with the main decoding results. These findings indicate that the PCu encodes force intensities regardless cue color and that above-chance decoding do not depend on whether betas were sorted by force level color or by the later-cued effector. The consistency of this control analysis with the main decoding results supports the conclusion that the PCu encodes force intensities in an abstract, effector-independent format during the first delay period. Second, label-permutation testing confirmed that the prediction accuracies obtained in the SVR analysis reflected the parametric coding of grip-force intensities. Specifically, computation of parametric contrasts on prediction accuracy maps derived from label-permuted models revealed highly similar clusters to those identified in the main analysis across all three time periods (at *p* < 0.05, FWE-corrected). Conversely, computation of t-contrasts on prediction accuracy maps reflecting the maximally permuted label showed no regions with above-chance decoding (even when tested at a liberal threshold of *p* < 0.001, uncorrected). Third, univariate analyses revealed no significant parametric modulation during the two delay periods and only in contralateral M1 during motor execution (even when tested at *p* < 0.001, uncorrected). This makes it unlikely that the parametric effects observed in the main MVPA analyses were primarily driven by univariate signal differences.

Taken together, these results, along with key features of our experimental design—including the orthogonality between visual features of the *Grip-force cue* and the motor information relative to the grip-force intensity, as well as the balanced presentation of visual stimuli (e.g., the frequency with which each grip-force level and the effector cues were presented in a given color)—indicate high specificity and sensitivity of the reported decoding effects.

## 5 CONCLUSION

This fMRI study provides new insights into how intended grip-force intensities are represented and transformed across subsequent stages of motor planning. Using a delayed grip-force task and time-resolved SVR, we found that parametric codes of intended grip-force intensities undergo a reformatting process, transitioning from abstract to effector- and movement-specific representations along the parieto-frontal axis. During the first delay, grip-force intensities were encoded in the PCu in an abstract, effector-independent format. In contrast, during the second delay, parametric activity patterns in the contralateral aIPS and PMd reflected effector- and movement-specific coding.

Furthermore, cross-effector and cross-regression decoding indicated that abstract representations in the precuneus are maintained across delays but not during execution, whereas effector-specific patterns in the aIPS and PMd closely resemble those observed during execution, reflecting movement-specific coding during the second delay period. These findings support a two-stage model of motor planning, in which abstract motor goals (i.e., intended force intensities) are progressively reformatted into effector-and movement-specific plans (i.e., grip-force anticipation) prior to execution.

In sum, our results indicate that the human brain encodes grip-force intensity through a structured sequence of transformations—from abstract codes to movement planning—across the parieto-frontal circuit. This process aligns with current models of motor planning and provides new evidence for a detailed and cohesive framework describing how abstract motor intentions are progressively transformed into concrete, executable movement plans. Future studies may investigate whether similar transformations occur across other effectors or motor domains.

## DATA AVAILABILITY STATEMENT

The data that support the findings of this study are available upon request from the corresponding author. The data are not publicly available due to privacy or ethical restrictions.

## ACKNOWLEDGMENT

This work was supported by DAAD: German Academic Exchange Service (https://www.daad.de/en/), Berlin School of Mind and Brain, Humboldt Universität zu Berlin (http://www.mind-and-brain.de/home/) and by the German Research Foundation (project grants to FB: Research Unit 5429/1) (467143400, BL 977/4-1).

## CONFLICT OF INTEREST

The authors have no known conflict of interest to declare.

## ETHICS APPROVAL STATEMENT

Participants gave their written consent before proceeding with the experiment. The experimental design and materials were approved by the ethics committee at Freie Universität Berlin (003/2021, Berlin, Germany).

## AUTHOR CONTRIBUTIONS

GC: Conceptualization, Data acquisition, Data curation, Methodology, Formal analysis, Writing-original draft. TTS: Conceptualization, Methodology, Supervision, Writing – review and editing. FB: Conceptualization, Methodology, Supervision, Writing – review and editing.

## Notes

### Competing Interest Statement

The authors have declared no competing interest.

